# Wnt signaling regulates ion channel expression to promote smooth muscle and cartilage formation in developing mouse trachea

**DOI:** 10.1101/2023.01.10.523309

**Authors:** Nicholas X. Russell, Kaulini Burra, Ronak Shah, Natalia Bottasso-Arias, Megha Mohanakrishnan, John Snowball, Harshavardhana H. Ediga, Satish K Madala, Debora Sinner

## Abstract

Ion channels play critical roles in the physiology and function of the nervous system and contractile tissue; however, their role in non-contractile tissue and embryonic development has yet to be understood. Tracheobronchomalacia (TBM) and complete tracheal rings (CTR) are disorders affecting the muscle and cartilage of the trachea and bronchi, whose etiology remains poorly understood. We demonstrated that trachealis muscle organization and polarity are disrupted after epithelial ablation of Wls, a cargo receptor critical for the Wnt signaling pathway, in developing trachea. The phenotype resembles the anomalous trachealis muscle observed after deletion of ion channel encoding genes in developing mouse trachea. We sought to investigate whether and how the deletion of *Wls* affects ion channels during tracheal development. We hypothesize that Wnt signaling influences the expression of ion channels to promote trachealis muscle cell assembly and patterning. Deleting *Wls* in developing trachea causes differential regulation of genes mediating actin binding, cytoskeleton organization, and potassium ion channel activity. Wnt signaling regulated expression of *Kcnj13, Kcnd3, Kcnj8,* and *Abcc9* as demonstrated by in vitro studies and in vivo analysis in *Wnt5a* and β-*catenin* deficient tracheas. Pharmacological inhibition of potassium ion channels and Wnt signaling impaired contractility of developing trachealis smooth muscle and formation of cartilaginous mesenchymal condensation. Thus, in mice, epithelial-induced Wnt/β-catenin signaling mediates trachealis muscle and cartilage development via modulation of ion channel expression, promoting trachealis muscle architecture, contractility, and cartilaginous extracellular matrix. In turn, ion channel activity may influence tracheal morphogenesis underlying TBM and CTR.

## INTRODUCTION

Ion channels play an essential role in the nervous system and in contractile tissues by controlling ion homeostasis and facilitating influx of ions mediating cell membrane potentials. While the physiological role of diverse ion channels has been studied in detail, their involvement in influencing tissue formation and patterning of non-contractile or non-nervous tissue is less well understood. An emergent area of research identifies channelopathies as underlying causes of craniofacial and other congenital defects. For example, the absence or lack of activity of the inwardly rectifying potassium ion channel KCNJ2 is associated with Andersen-Tawil syndrome. Loss of function of KCNJ2 alters bioelectric signaling (Adams et al. 2016) and Bmp/Smad1/3/5 signaling (Belus et al. 2018) thus impairing the patterning of the craniofacial tissue.

In the lung epithelium, ion channels play an essential role in regulation of fluid and periciliary functions, i.e. pH, viscosity and composition to control muco-ciliary clearance, achieved by the coordinated activity of ion channels CFTR, CaCC, ClC2 and ENaC (Bartoszewski, Matalon, and Collawn 2017). Potassium ion channels play roles in the maintenance of electrochemical gradients and ion homeostasis; thus, being necessary for chloride secretion and sodium reabsorption. ATP-dependent potassium channels, such as KCNJ8, link the metabolic status of the alveolar cells to the membrane potential which may be of relevance to the process of airway epithelial repair (Buchanan et al. 2013; Trinh et al. 2007).The activity of potassium channels may also be regulated by the O_2_ tension in epithelial cells of the lung. While the mechanism by which changes in O2 tension affect potassium channel activity has not been established, pharmacological modulation of potassium channels may be considered as a therapy to treat conditions associated with hypoxia (Bardou, Trinh, and Brochiero 2009). Since potassium channels regulate the membrane potential of smooth muscle cells, and thus the cytoplasmic free Ca^2+^ concentration (Jackson 2005), abnormal function of potassium channels has been linked to pulmonary hypertension(Redel-Traub et al. 2022). Potassium channels affect the actin cytoskeleton of developing trachealis smooth muscle via the AKT signaling pathway (Yin et al. 2018). In turn, the actin cytoskeleton is essential for the detection and transduction of mechanical stress to ion channels, inducing their activity (Martinac 2014).

In mice and humans, the large airways of the respiratory tract are patterned in the dorsal-ventral axis with trachealis muscle occupying the dorsal side, and cartilage rings spanning the ventro-lateral sides. Disrupting of dorsal-ventral patterning is associated with Tracheomalacia, a congenital disorder resulting in flaccid airways, or complete tracheal rings (CTR) wherein the muscle is absent, and the luminal surface of the airway is narrow (Sinner et al. 2019). Our previous studies have established that Wls-mediated Wnt signaling pathway plays an important role in the morphogenesis of the trachea affecting the formation of cartilage and the patterning of the trachealis muscle (Snowball et al. 2015). Wls, a cargo receptor that mediates Wnt ligand secretion, is essential for the activation of both the canonical pathway, which is mediated by β-catenin, and the non-canonical Wnt signaling pathway, and plays essential roles in respiratory tract development (Banziger et al. 2006; Bartscherer et al. 2006; Cornett et al. 2013; Jiang et al. 2013).

The Planar Cell Polarity (PCP) branch of the non-canonical Wnt signaling pathway, plays important roles in the normal organization and functioning of the trachealis smooth muscle (Kishimoto et al. 2018). Previous studies determined that ion channels, particularly the calcium-activated chloride channels and voltage-gated potassium channels, play an important role in determining smooth muscle cell shape and organization. In mice, deletion of *Tmem16 (Ano1)* results in abnormal organization of the smooth muscle of developing trachea, and affects the formation of the tracheal cartilage (Rock, Futtner, and Harfe 2008). Kcnj13, an inwardly rectifying potassium ion channel, is critical for trachealis muscle organization, tubulogenesis, and lung development (Villanueva et al. 2015; Yin et al. 2018). While compelling evidence indicates a role of ion channels in trachealis muscle and cartilage patterning(Lin et al. 2014; Bonvin et al. 2008; Wallace et al. 2013), mechanisms underlying the process remain unclear.

In the present study we determined that Wnt/β-catenin mediated signaling is essential for smooth muscle patterning and assembly and for regulation of ion channel expression during tracheal development. In mice, abnormal activity of inwardly rectifying ion channels impaired trachealis smooth muscle and cartilage development. Thus, a tight control of ion channels expression and activity during development of the trachea is necessary for the proper patterning of the large airways.

## MATERIAL AND METHODS

### Mouse breeding and genotyping

Animals were housed in a pathogen-free environment and handled according to the protocols approved by CCHMC Institutional Animal Care and Use Committee (Cincinnati, OH, USA). Generation of the *Wntless* (*Wls*) conditional knock-out (CKO) mouse has been described (Carpenter et al. 2010). *Shh^Cre/wt^ ; Wls^f/f^* embryos were obtained by breeding *Wls^f/f^* mice with *Shh^Cre/wt^* mice and rebreeding the resulting mice with *Wls^f/f^* mice (Harfe et al. 2004). Generation of *γSMA eGFP* mice was previously described (Szucsik et al. 2004) *ShhCre;Wls^f/f^;γSMAeGFP* embryos were generated by breeding *γSMAeGFP* mice with *ShhCre;Wls ^f/wt^*mice and rebreeding the resulting mice with *γSMAeGFP*;*Wls^f/f^* mice. *β-catenin^f/f^*mice (Jackson Laboratories, strain 004152 (Brault et al. 2001) mice were mated with *Foxg1^Cre/wt^* to generate *Foxg1^Cre/wt^*;*β-catenin^f/wt^*. Embryos of genotype *Foxg1^Cre/wt^*;*β-catenin^f/f^ were generated by mating β-catenin^f/f^ with Foxg1^Cre/wt^;β-catenin^f/wt^. Wnt5a^f/f^* were mated with *Dermo1^Cre/wt^* to generate *Dermo1^Cre/wt^;Wnt5a^f/wt^* mice and rebreed to with *Wnt5a^f/f^* generating embryos of genotype Dermo1Cre;Wnt5a*^f/f^* mice ((Ryu et al. 2013), Jackson laboratories strain # 026626). *Ror2^f/f^* (Jackson laboratories strain # 018354 (Ho et al. 2012)) mice were mated with *Foxg1^Cre/wt^*and the resulting mice rebreed with *Ror2^f/f^* to generate embryos of genotype *Foxg1^Cre/wt^; Ror2^f/f^*. *Sox9KIeGFP* mice was previously described ((Chan et al. 2011) Jackson laboratories strain 030137). Genotypes of transgenic mice were determined by PCR using genomic DNA isolated from mouse-tails or embryonic tissue. Primers utilized for genotyping are provided as Supplementary material (Supplementary table 1).

### Transcriptomic analyses

RNA-sequencing data from E13.5 *ShhCre;Wls^f/f^* tracheas vs E13.5 *Wls^f/f^*tracheas were obtained from our previously published work ((Bottasso-Arias et al. 2022) GEO repository under GSE158452). Differentially expressed genes were identified using Deseq (N=5 Controls, N=4 Mutants) (Anders and Huber 2010; Lawrence et al. 2013). Fragments Per Kilobase (FPKM) values were calculated using Cufflinks (Trapnell et al. 2010). Differentially expressed genes had a p-value <.05, FC>1.5 and FPKM>1 in over half of the replicates in at least one condition. The same approach was utilized for gene expression analysis of control chondroblast, epithelium and smooth muscle cells isolated by FACS. (GEO repository GSE241175). Heatmaps were generated using normalized counts generated by DEseq and pheatmap or from RNA-seq fold changes. Functional enrichment was performed using Toppfun and hits relevant to this project were visualized in a -log10 (pvalue) bar graph. System models were created using IPA’s Path Designer ("IPA").

<u>Phalloidin staining: Tissue was fixed in 4% PFA and embedded in OCT to generate 7μm sections. Staining was performed as recommended by the supplier with the following modifications. Slides were equilibrated at room temperature and rinsed in PBS twice, and permeabilized for 15 minutes in 0.1% Triton/1XPBS The slides were then rinsed in 1XPBS, and incubated for 30 minutes in Alexa Fluor 488 Phalloidin (Invitrogen A12379), at 1:40 diluted accordingly in the blocking solution 1. Following incubation slides were rinsed in PBS and cover-slipped with Prolong Gold containing DAPI.</u>

### Immunofluorescence staining

Embryonic tissue was fixed in 4% PFA overnight and embedded in paraffin or OCT to generate 7μm sections. For general immunofluorescence staining, antigen retrieval was performed using 10mM Citrate buffer, pH6. Slides were blocked for 2 hours in 1XTBS with 10% Normal Donkey serum and 1% BSA, followed by overnight incubation at 4^0^C in the primary antibody, diluted accordingly in blocking solution. Slides were washed in 1X TBS-Tween20 and incubated in secondary antibody at 1:200, diluted accordingly in blocking solution, at room temperature for one hour and were then washed and cover-slipped using Vecta shield mounting media with or without DAPI. Fluorescent staining was visualized and photographed using automated fluorescence microscopes (Zeiss and Nikon). Antibodies utilized in this manuscript have been previously used and validated by our laboratory and other investigators. Source, references, and dilution of primary and secondary antibodies used have been provided as Supplementary material (Supplementary table 2).

### Whole mount staining

Tracheal lung tissue isolated at E13.5 was subject to whole mount immunofluorescence as previously described (Sinner et al. 2019). Embryonic tissue was fixed in 4% PFA overnight and then stored in 100% Methanol (MeOH) at - 20^0^C. For staining, wholemounts were permeabilized in Dent’s Bleach (4:1:1 MeOH: DMSO: 30%H_2_O_2_) for 2 hours, then taken from 100% MeOH to 100% PBS through a series of washes. Following washes, wholemounts were blocked in a 5% BSA (w/v) blocking solution for two hours and then incubated, overnight, at 4^0^C in primary antibody diluted accordingly in the blocking solution. After 5 one-hour washes in PBS, wholemounts were incubated with a secondary antibody at a dilution of 1:500 overnight at 4^0^C. Samples were then washed three times in 1X PBS, transferred to methanol, and cleared in Murray’s Clear. Images of wholemounts were obtained using confocal microscopy (Nikon A1R). Imaris imaging software was used to convert z-stack image slices obtained using confocal microscopy to 3D renderings of wholemount samples.

### Tracheal mesenchymal cell Isolation and culture

Primary cells were isolated as previously described (Gerhardt et al. 2018). Briefly, E13.5 tracheas of at least five embryos of the same genotype were isolated, washed in 1X PBS, minced in TrypLE express (Gibco) and incubated for 10 minutes at 37^0^C. After incubation, tissue was pipetted until cell suspension formed. Cells were seeded in flasks containing MEF tissue culture media composed of DMEM, 1% penicillin/streptomycin, 2% antibiotic/antimycotic, and 20% non-heat inactivated FBS. Only mesenchymal cells were attached, as we confirmed expression of *Sox9, Col2a1* and *Myh11* but no expression of *Nkx2.1* was detected (data not shown). For studies involving monolayers, 1x10^5^ cells were seeded in 6-well plates. Some monolayers were pre-treated with DMSO (vehicle, ICN19141880 Fisher), Minoxidil sulfate (50 μM; M7920, Sigma) and VU590 (15-25 μM; 3891,Tocris) for 48 hours, with treatments being change every 24 hours. These cells were subsequently used in the following micromass assays. For micromasses experiments, 5x10^4^ pre-treated cells in 8μl drops were seeded per well in a 8-well slide. and flooded with culture media after 2 hours. All micromasses were treated with rBMP4 (250-500 ng/mL; 314-BP-050, R&D). As specified in the results section, after the micromasses were seeded, some of the DMSO pre-treated cells received DMSO (vehicle), Minoxidil sulfate (50 μM), or VU590 (15-25 μM) as a post-treatment. All cells were treated on the consecutive day and harvested after a week, with treatments changed every 48 hours. Alcian Blue staining was performed to evaluate production and secretion of glycosaminoglycans into the extracellular matrix. Images were obtained using a Nikon wide field microscope coupled with a DS-Fi3 color camera.

### Dissociation and FACS of embryonic trachea

Cell dissociation was performed following a previously published protocol (Bottasso-Arias et al. 2022). E13.5 γSMAeGFP tracheas were dissected in cold PBS and dissociated to single cells using TrypLE Express (phenol-red free, Thermo, 12604013) at 37°C for 10 minutes, followed by trituration for 30 seconds at RT. Cells were washed twice with FACS buffer (1mM EDTA, 2% FBS, 25mM HEPES in phenol-red free HBSS). To identify epithelial cells, cells were stained with APC anti-mouse CD326/EpCAM (Invitrogen, ref 17-5791-82, used at 1:50) at 4°C for 30 minutes followed by two washes with FACS buffer. Cells were resuspended in FACS buffer and passed through a 35 µm cell strainer. To stain dead cells, Sytox Blue nucleic acid stain (Thermo, S11348, used at 1 µM) was added to the cell suspension. Cells were sorted using a BD FACS Aria I and II. Single live “chondroblast” cells were collected after size selection and gating for Sytox-negative, EpCAM-negative, and eGFP negative cells. Cells were sorted directly into RNA lysis buffer (Zymo Research Quick RNA Micro kit, S1050) for isolation of RNA.

### Embryonic tracheal-lung culture

Embryonic tracheas were harvested at E11.5 or E12.5 and cultured at air-liquid interphase as described (Hyatt, Shangguan, and Shannon 2004). DMSO, WntC59 (15μM, 5148 TOCRIS), VU590 (25 μM), and Minoxidil Sulfate (50 μM) were added to the culture one hour after setting the tissue in the inserts. Photographs or videos of the tissue were taken every 24 hrs. After 72 hours the tissue was harvested and processed for paraffin embedding. Culture media and treatments were replaced every 48 hours.

### In-Situ Hybridization

The procedure was performed according to a protocol developed by Advanced Cell Diagnostics (ACD) (Wang et al. 2012). In situ probes were designed by ACD. Slides were baked and deparaffinized. In situ probes were added to the slides and hybridization was performed for 2 hours at 40^0^C, followed by several rounds of amplification steps. For fluorescence detection, opal dyes were utilized to detect the localization of the transcripts. After mounting with permanent mounting media, slides were photographed using a wide field Nikon fluorescent microscope.

### RNA isolation and RT-PCR

Gene expression was determined by quantitative RT-PCR. RNA was isolated from E13.5 tracheas and lungs using commercially available kit (Zymo Research Quick RNA Micro kit, S1050). Reverse transcription was performed according to manufacturer instructions (High-Capacity cDNA Reverse Transcription kit, Applied Biosystems), and Taqman probes were utilized to detect differential expression using StepOne Plus and QuantStudio3 RT-PCR system. Gene expression was normalized to *Gapdh*.

### Motif Enrichment Analyses

Promoter regions 2Kb upstream of the predicted transcriptional start site of select genes were analyzed using MEME suite’s FIMO coupled with Meme Suite’s Motif database (Grant, Bailey, and Noble 2011). Promoter sequences were downloaded using the UCSC Table browser. Selective motif locations in DNA sequences of interest were identified using JASPAR. IGV was used to visualize motif and luciferase construct locations within promoters.

### Collagen Gel Assay

Free floating collagen gel assay was performed as previously described (Gajjala et al. 2021). In brief, 200,000 cells were seeded into rat tail collagen (1mg/ml; A10483-01, Gibco) gel matrices. Each well contained 500 μL consisting of 2/3 volume cells resuspended in MEF media culture (see Tracheal mesenchymal cell isolation and culture section) and 1/3 volume of rat collagen. For each well, 5 μL of 1M NaOH was also added to initiate collagen gel polymerization and the mixture plated in a 24 well plate. After 15 minutes, the solidified gels were detached from the walls of the well using the tip of a pipette. Each well was filled with 1 mL of cell media and treatments consisting of DMSO, WntC59 (15μM), JNK inhibitor II (20μM, 420119 Millipore Sigma), VU590 (25μM), XAV939 (50μM, 3748 TOCRIS), Diazoxide, or Minoxidil Sulfate (25μM and 50μM) were added. Collagen gel images were taken every six to twelve hours and the surface area of each gel was determined using EVOS analysis software. The areas acquired during each timepoint were compared to the original area of the gel to determine the magnitude of its contraction and plotted as the percentage of contraction vs time.

### Videos

Time lapse imaging to visualize contraction was obtained using EVOS M7000. Tracheal lung tissue was cultured in ALI with treatments as described in Tracheal-lung culture section. Imaging was obtained over a period of 30 seconds at room temperature. These pictures were assembled to form a video in which duration and amplitude of contraction could be determined for differently treated tracheal samples.

### Statistics

Quantitative data were presented as mean ± standard error. For animal experiments, a minimum of three different litters for each genotype were studied. Experiments were repeated at least twice with a minimum of three biological replicates for each group. Statistical analysis was performed using Graph Pad Prism ver.8.2.0 and 9.0.0 for MacOS. Statistically significant differences were determined by paired T-test, or one-way or two-way ANOVA repeated measures followed by post hoc pairwise multiple comparison procedures (Dunnet or Holm-Sidak test). Significance was set at P<0.05.

## RESULTS

### Canonical and non-canonical Wnt signaling play distinct roles in smooth muscle cell organization of the developing trachea

Epithelial deletion of *Wls* causes a fundamental disruption to the patterning of the trachealis muscle being ectopically localized, and oriented parallel to the elongation axis of the trachea as opposed to the transverse organization observed in the control developing trachea (Fig1A,B)(Snowball et al. 2015). The changes in muscle cell organization and trachealis muscle assembly indicate anomalies in cytoskeletal organization of the trachealis muscle cells of *ShhCre;Wls^f/f^* samples. RNA seq. analysis demonstrated functional enrichment and induction of genes supporting the actin cytoskeleton in *Wls* deficient tracheas (Fig1C). RNA seq data was consistent with the morphological changes observed in the trachealis muscle of the *ShhCre;Wls ^f/f^* embryos, wherein anomalous muscle cell shape was detected by actin smooth muscle antibody and phalloidin staining (Fig1A,B SFig1). To identify the localization of Wnt ligands that may be affected by deletion of *Wls*, we performed RNA in situ hybridization on E13.5 transverse sections of developing trachea. *Wnt7b* transcripts were primarily detected in tracheal epithelium, while *Wnt5a* transcripts were enriched in the ventral tracheal mesenchyme. *Wnt4* was detected in the epithelium enriched in the dorsal aspect (Fig1D). Since *Wls* is required for secretion of ligands triggering both canonical and non-canonical Wnt signaling, and β-catenin independent Wnt signaling is associated with changes in cytoskeletal organization, we compared the phenotype observed in *ShhCre;Wls^f/f^* trachea to the phenotypes observed after mesenchymal deletion of *β-catenin* or *Wnt5a* in developing tracheas. While the organization of the muscle after ablation of *Wnt5a* in tracheal mesenchyme of developing embryo is characterized by cytoskeletal changes resulting in lack of alignment and orientation among the muscle cells, evident in the proximal region, the localization of the trachealis muscle remained confined to the dorsal aspect of the trachea (Fig1E,F). In agreement with previous studies, the *Wnt5a* deficient tracheal mesenchyme appears thicker than in control due to the increased number of layers of Sox9+ cells detected in the ventro-lateral aspect of the trachea (Fig1F). Similarly, mesenchymal deletion of *Ror2*, the receptor of Wnt5a, altered the organization of the smooth muscle cells without changing the dorsal localization of the trachealis muscle (Fig1F). On the other hand, lack of mesenchymal β-catenin in *Foxg1Cre;β-catenin^f/f^* caused a drastic reduction of Sox9+ cells and an unexpected profound effect on the organization of the muscle cells. This abnormal organization resembled the *Wls* deficient trachea wherein myocytes were ectopically located in the ventral mesenchyme where chondroblasts are normally located (Fig1E, F). Taken together, epithelial Wls-induced Wnt signaling is essential for the morphology and patterning of the trachealis muscle, primarily acting via Wnt/β-catenin signaling likely induced by epithelial Wnt7b. Disruption of Wnt5a-Ror2-mediated signaling impairs the organization of the smooth muscle without having a major impact on the dorsal-ventral fate of the mesenchymal cells.

**Figure1:**
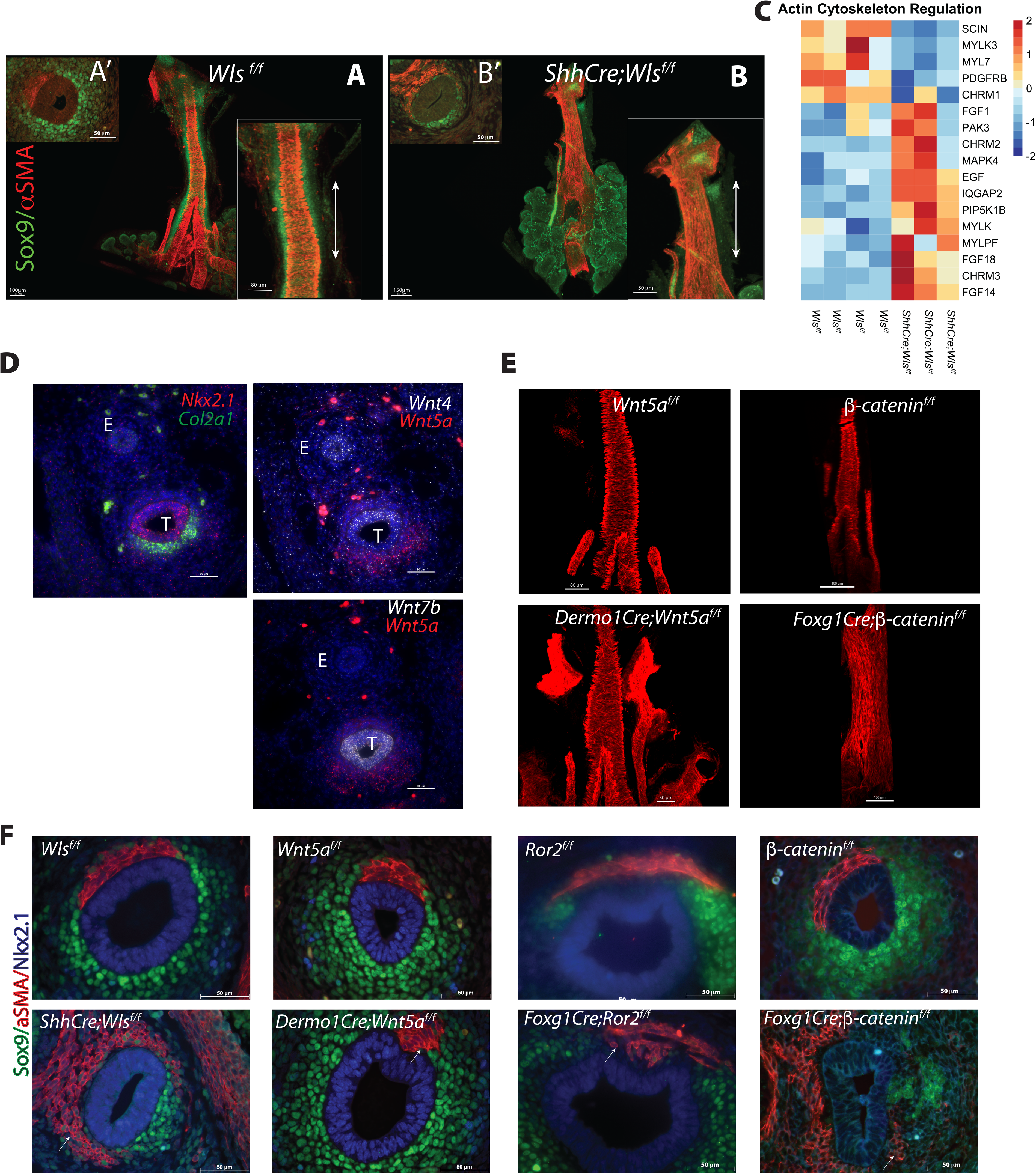
Canonical and Non-canonical Wnt signaling play distinct roles in trachealis smooth muscle cell patterning and organization. A) Whole mount images of E13.5 *Wls^f/f^* (control) trachea-lung are shown, inset depicts a higher magnification of the dorsal view of the trachea. A’) Cross section of an E13.5 control trachea demonstrating ventral localization of chondroblasts (Sox9+ cells) and dorsal localization of trachealis muscle cells. B) Epithelial deletion of *Wls* disrupts tracheal smooth muscle cell organization and morphology. In *ShhCre;Wls^f/f^* tracheas, muscle is ectopic and oriented parallel to the tracheal elongation axis (Compare insets in A and B). Sox9 is seldom detected on *Wls* deficient tracheal mesenchyme as determined by immunofluorescence of cross section (B’) and whole mount tissue staining. Double-headed arrows in insets A’ and B’ indicate orientation of the tracheal elongation axis. C) Heat map depicting changes in expression of genes influencing cytoskeleton organization detected in E13.5 *Wls* deficient tracheas. D) RNA in situ hybridization depicts localization of transcripts for Nkx2.1, Col2a1, Wnt4, Wnt5a and Wnt7b in transverse tracheas at E13.5. Note the epithelial localization of Wnt7b and Nkx2.1. Wnt4 transcripts are enriched are primarily observed to the dorsal aspect of the tracheal epithelium, while Wnt5a transcripts are observed in the ventral tracheal mesenchyme overlapping with the localization of Col2a1. E) Whole mount stainings of E13.5 tracheas depicting the abnormal organization of the smooth muscle after deletion of *Wnt5a* and *β-catenin* in tracheal mesenchyme. Note that after deletion of β-catenin smooth muscle cells are ectopic and abnormally oriented parallel to the elongation axis reminiscent of the *Wls* deficient phenotype F) Cross section staining of E13.5 tracheas depicting the abnormal shape, patterning, and lack of organization of the trachealis smooth muscle cells after deletion of *Wls, Wnt5a*, *Ror2,* or *β-catenin*. While *Wnt5a* or *Ror2* deficient tracheas have altered shape and assembly of smooth muscle cells, only deletion of *Wls* and *β-catenin* caused ectopic localization of smooth muscle.

### Epithelial deletion of *Wls* causes differential expression of potassium ion channels in developing trachea

The lack of mesenchymal organization in the *ShhCre;Wls^f/f^*mouse caused by lack of β-catenin signaling, presumably alters the expression of molecules affecting smooth muscle organization and function. While performing Gene ontology analysis, we identified an enrichment of molecular functions linked to ion channel activity in the embryonic *ShhCre;Wls^f/f^* trachea. Those genes encode potassium ion channels such as KCNJ13, KCNJ8, as well as the enzyme PRSS8 necessary for activation of the ENaC channel and the chloride channel CFTR (Figure2A). Functional enrichment analysis identified pathways associated with the differentially upregulated ion channels being involved in muscle and cardiac contraction, and potassium ion channels (Fig2B). Pathways enriched among differentially downregulated ion channels included voltage gated potassium ion channels as well as stimuli-sensing ion channels (Fig2B). To validate the RNAseq data, we examined the expression pattern of several ion channels predicted to be differentially regulated in *ShhCre;Wls^f/f^* (1.5-fold change expression) and for which a potential role or association with respiratory tract homeostasis and disease has been described. We focused on *Kcnc2, Kcnd3, Kcnk3, Kcnip1, Kcnj13, Kcnj8*, and its related subunit *Abcc9*. *Ano1,* which enables intracellular calcium activated chloride channel activity and is known to affect tracheal morphogenesis and its paralog, *Ano4*, were also included in the analysis. We examined their normal expression pattern by FACS of E13.5 control tracheal tissue (Fig2C) and RNA in situ hybridization (Fig2D) and observed that while some ion channels were detected across the tracheal tissue, several ion channels were enriched in specific compartments of the trachea. *Kcnc2 and Ano4* were enriched in the ventral aspect of the trachea where cartilage forms. *Ano1, Kcnd3,* and *Kcnip1(not shown)* were enriched in tracheal myocytes. *Cftr* and *Kcnj13* transcripts were enriched in tracheal epithelium. On the other hand, the ion channel *Kcnj8 and its gating subunit Abcc9*, were expressed in the muscle of the developing esophagus whereas their expression in normal developing trachea was low and barely detected by RNA in situ hybridization (Fig2D). Ion channels differentially regulated in *Wls* deficient trachea were also expressed in developing lung; however, some of these genes were enriched in the trachea such as *Kcnc2* and *Kcnip1*; while *Ano4* was more abundant in developing lung as determined by qRT-PCR of tracheal and lung tissue (Fig2E). Taken together, ion channels which were differentially regulated by Wnt signaling were expressed in developing tracheal tissue with unique and overlapping expression patterns.

**Figure2:**
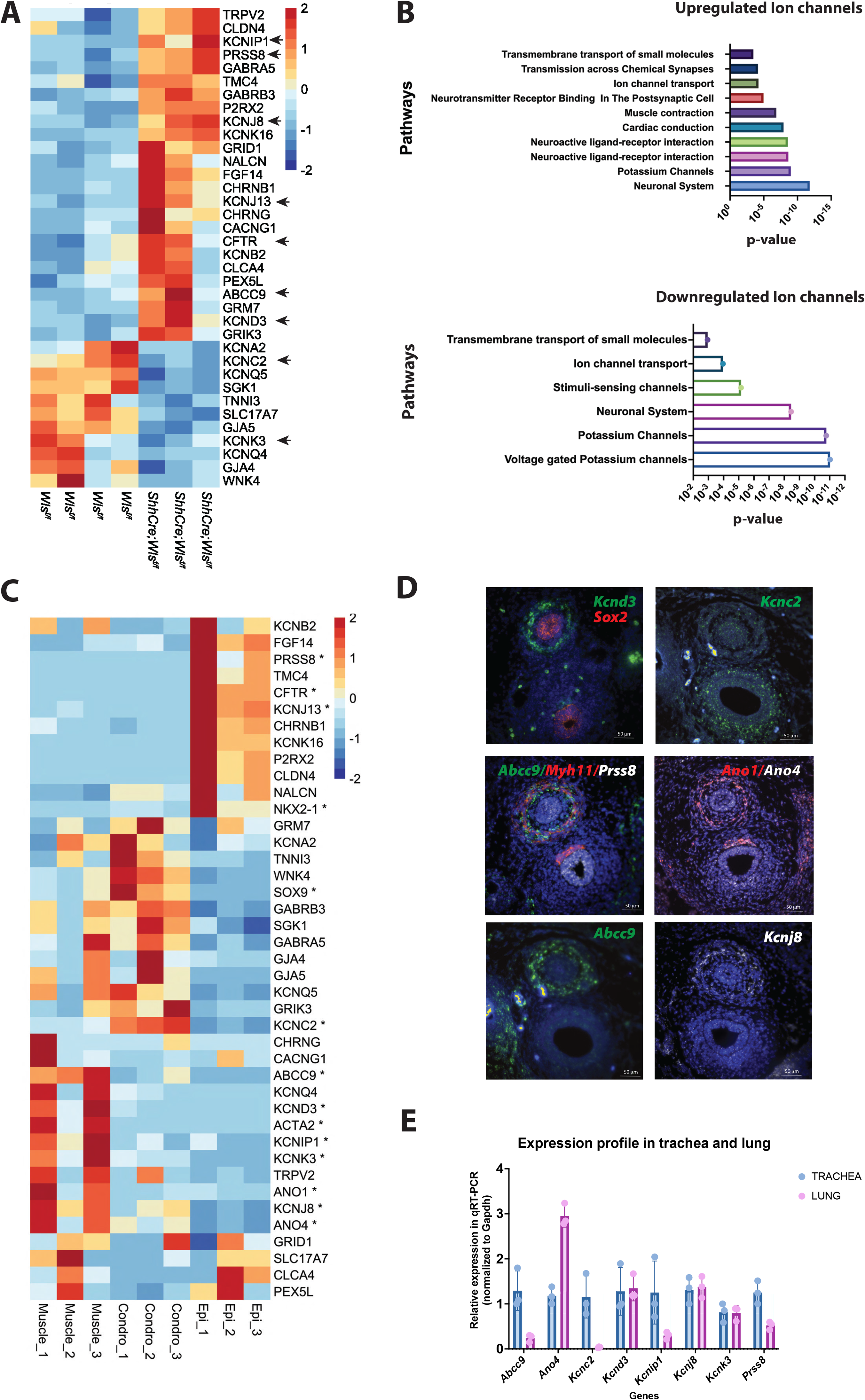
Epithelial deletion of *Wls* alters expression of potassium ion channels in developing trachea. A) Heat map depicting potassium ion channel encoding genes differentially regulated in response to Wnt signaling. B) GO term enrichment analysis identified pathways associated with ion channels upregulated or downregulated after the deletion of *Wls* in embryonic trachea. C) Heat map of ion channel encoding genes demonstrates differential expression patterns among epithelial, muscle, and cartilaginous cells of the trachea at E13.5. Expression of *Nkx2.1, Sox9*, and *Acta2* confirmed the epithelial, cartilaginous, and muscle identity of the cell isolation respectively. *Kcnc2* is expressed in chondrogenic cells, *Kcnj8* and *Abcc9* are present in trachealis muscle cells, while *Kcnj13* and *Cftr* are expressed in epithelial cells. D) RNA scope in situ hybridization depicting differential localization and expression of ion channel transcripts and related molecules in epithelium or mesenchyme of E13.5 esophagus and trachea. E) RT-PCR analysis demonstrates differential levels of transcripts in tracheas and lungs at E13.5. *Abcc9, Kcnc2, Kcnip1*, and *Prss8* are relatively more abundant in trachea than lungs, while *Ano4* is more abundant in lungs. N=3.

### Decreased Wnt-β-catenin dependent activity is associated with increased of potassium ion channel expression

After determining the expression pattern of genes predicted to be differentially regulated by the epithelial deletion of *Wls*, we proceeded to analyze their expression patterns in the Wls deficient tracheal tissue. qRT-PCR and RNA in situ hybridization studies confirmed the robustness of the RNAseq data. *Ano4* and *Kcnc2* were decreased while *Kcnd3*, *Kcnj8*, and *Abcc9* were increased and ectopically expressed in trachea after deletion of *Wls. Prss8 RNA* was increased in the epithelium of *Wls* deficient tracheas*. Notum,* a direct target of Wnt/β-catenin was decreased (Fig3A, B). To define the mechanism by which the epithelium modulates ion channel expression in developing trachea downstream of *Wls*, we tested whether mesenchymal deletion of *β-catenin* affected the expression of ion channels differentially regulated by *Wls*. Loss of mesenchymal β-catenin caused similar changes in the expression of ion channels as observed in *Wls* deficient tracheas. *Kcnj8* and its regulatory unit *Abcc9* and *Kcnd3* were upregulated while *Kcnc2* trended downregulated after mesenchymal deletion of β-catenin. *Notum,* a target of Wnt/β-catenin, was downregulated after mesenchymal deletion of β-catenin (Fig.3C, D). Despite the changes observed in muscle cell cytoskeleton, expression of these genes encoding ion channels were unchanged in the *Dermo1Cre;Wnt5a^f/f^* mutant tracheas (Fig3C). Since mesenchymal deletion of *β-catenin* causes increased expression of muscle markers, we tested whether the changes observed in expression of *Kcnj8* and *Abcc9* were due to the higher number of smooth muscle cells present in β-catenin deficient trachea and normalized the levels of *Kcnj8* in relation to *Myocd*. RNA in situ hybridization and qRT-PCR studies demonstrated that despite the increased expression of *Myocd* or *Acta2* in *Foxg1Cre;β-catenin^f/f^* tracheas the levels of *Kcnj8* were augmented after normalization with the muscle markers. Next, we analyzed the 2kb promoter regions of human *KCNJ8* and *ABCC9* and identified several potential binding sites for LEF1 and TCF3/4 (SFig2) suggesting that in mouse developing trachea *Wls* via β-catenin may modulate the expression of *Kcnj8* and *Abcc9* by binding to regulatory sequences. However, no LEF1 or TCF3/4 binding sites were detected on the *KCND3* promoter. Thus, epithelial-induced-Wnt signaling influences the actin cytoskeleton and modulates expression of *Kcnd3, Kcnj8, and Abcc9* in developing trachea primarily via β-catenin.

**Figure3:**
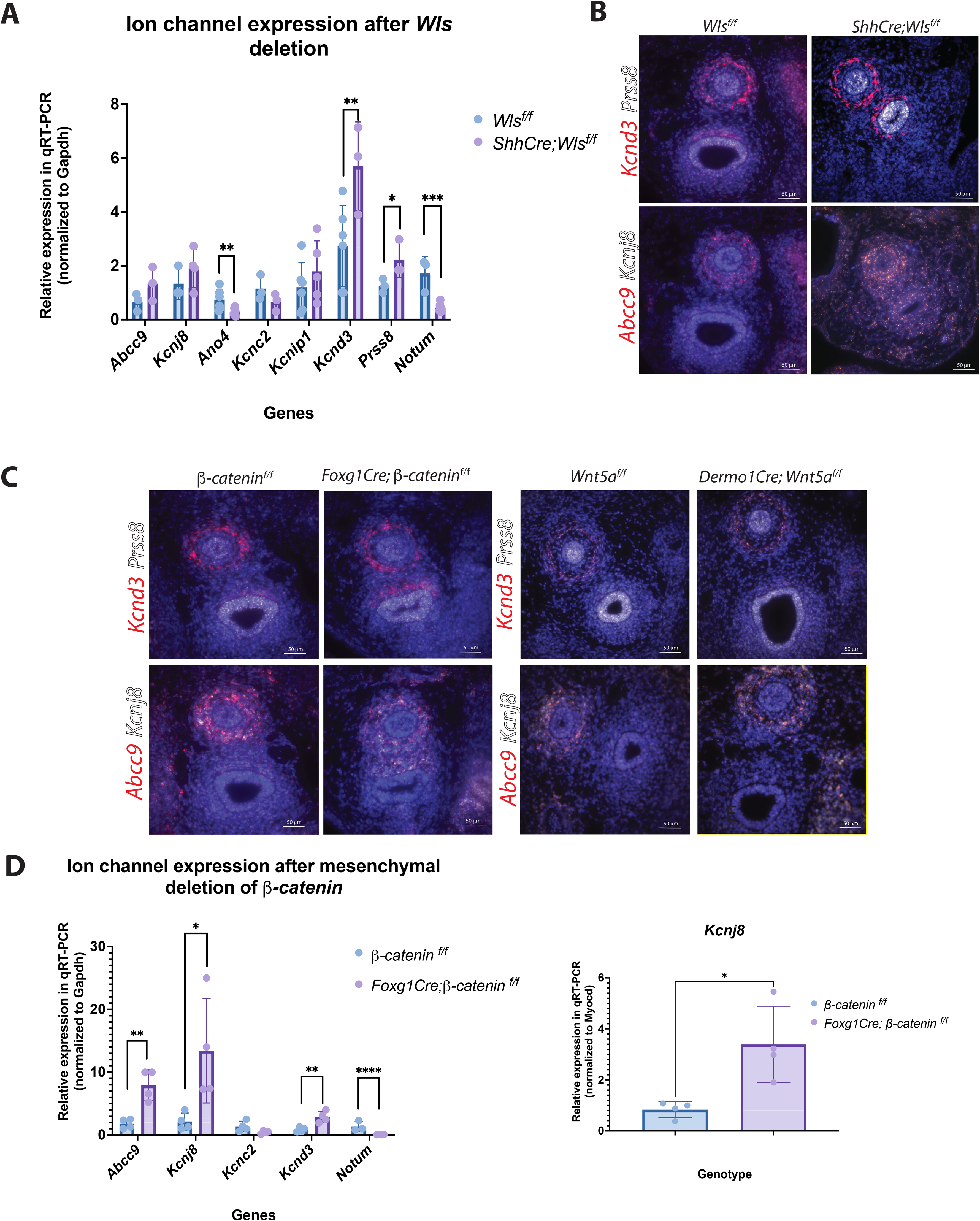
Decreased Wnt/β-catenin activity is associated with increased potassium ion channel expression. A) RT-PCR performed on E13.5 *Wls^f/f^* and *ShhCre;Wls^f/f^*tracheas detected decreased expression of *Ano4* and *Notum* (a direct target of Wnt/β-catenin), and increased *Kcnd3* expression. T-test: *p<0.05 **p<0.01 N=5. B) RNA scope in situ hybridization on E13.5 tracheal cross sections depicting localization of *Prss8* in *Wls^f/f^* tracheal epithelium. *Kcnd3, Kcnj8*, and *Abcc9* are strongly detected in mesenchyme surrounding the esophagus and at lower levels in the tracheal tissue. In *ShhCre;Wls^f/f^*trachea, *Prss8, Abcc9, Kcnj8, and Kcnd3* RNA levels were increased. *Abcc9* and *Kcnj8* transcripts were ectopically located in the tracheal mesenchyme and epithelium. C) RNA scope in situ hybridization demonstrates increased *Kcnd3, Prss8, Abcc9*, and *Kcjn8* RNAs in *β-catenin* deficient tracheas; Mesenchymal deletion of *Wnt5a* did not affect expression of these genes. D) RT-PCR analysis performed on E13.5 *Foxg1Cre;β-catenin^f/f^* tracheas support the RNA scope findings. *Abcc9, Kcnj8*, and *Kcnd3* are increased after mesenchymal deletion of β-*catenin*, while *Notum* is downregulated. Despite the increased number of smooth muscle cells, *Kcnj8* RNA was increased in *Foxg1Cre;β-catenin ^f/f^* trachea as determined after normalizing *Kcnj8* transcript levels to *Myocd*. T-test: *p<0.05 **p<0.01, ****p<0.001 N=5

### Abnormal ion channel activity and deficient Wnt signaling activity impair the organization and contractility of the developing trachealis muscle ex vivo

Analysis of RNA seq data indicated that differentially regulated ion channels may influence smooth muscle contractility after the deletion of *Wls* (Fig4A). To determine whether myocytes in Wls-deficient trachea were contractile despite their anomalous alignment, we performed in vivo studies wherein tracheal-lung tissue isolated from E12.5 *ShhCre;Wls^f/f^;γSMAeGFP* tracheas was cultured in air-liquid-interface (ALI) for 72 hr. In γSMAeGFP tissue contraction occurred perpendicular to the elongation axis of the trachea. On the contrary no contraction was detected in *ShhCre;Wls^f/f^;γSMAeGFP* tracheal tissue (SVideo1,2). Myocytes present in the bronchi contracted to some extent; however, the orientation of the contraction occurred parallel to the elongation axis of the trachea. This finding is distinct from control bronchi wherein muscle contraction occurs transverse to the longitudinal axis of the bronchi (SVideo2). Similarly, using primary mesenchymal cells isolated from *Wls^f/f^* and *ShhCre;Wls^f/f^*tracheas in a free floating collagen assay, we demonstrated that contraction of *Wls* deficient mesenchymal cells was impaired as indicated by the decreased contraction of collagen gels (Fig4B). These findings were reproduced in vitro in free-floating collagen assay wherein reduced contractility was observed when collagen gels containing *Wls^f/f^* cells were treated with the Wnt signaling inhibitor WNTC59 (Proffitt et al. 2013) (Fig4C). Treatment of cells with non-canonical Wnt inhibitor JNK inhibitor II (Bennett et al. 2001) or with β-catenin dependent Wnt signaling inhibitor XAV939 (Huang et al. 2009), did not have a pronounced impact on contractility (Fig4C and not shown). Activation of Kcnj8 with diazoxide or minoxidil sulfate (Gopel et al. 2000) did not have a strong effect on contractility as measured by the collagen assay, while WntC59 treatment impaired the contractility (Fig 4D). Since abnormal expression and activity of ion channels impaired contractility and the structure of the trachealis muscle, we sought to test in an *in vitro* system whether altered activity of selected ion channels may affect the organization and contraction of smooth muscle cells. Trachea-lung tissue from *γSMAeGFP* mouse was isolated at E12.5 and cultured in ALI for 72 hrs. Samples treated with DMSO (control), depicted the normal orientation of trachealis muscle fibers from time 0 to 72hrs. In tissue treated with VU590 (a blocker of KCNJ13 ion channel (Lewis et al. 2009)), there was a significant loss of trachealis muscle organization, as previously described (Yin et al. 2018). Treatment with WntC59 (Proffitt et al. 2013), caused disorganization of muscle cells an effect that was similar to the findings after the treatment with VU590 (Lewis et al. 2009)(Fig.4E). Activation of KCNJ8 with minoxidil sulfate or diazoxide did not affect trachealis muscle organization in ALI (Fig4E). Remarkably, blocking inwardly rectifying ion channels utilizing the pharmacological reagent VU590 (Lewis et al. 2009) severely impaired the contractility of the trachealis muscle cells ex vivo in ALI culture (SFig.3) while treatment with Minoxidil Sulfate did not impact contractility, which was similar to the findings from the collagen assay (SFig.3). Taken together, the data support a role for Wnt signaling and potassium ion channels influencing the organization and contractility of the developing trachealis muscle.

**Figure4:**
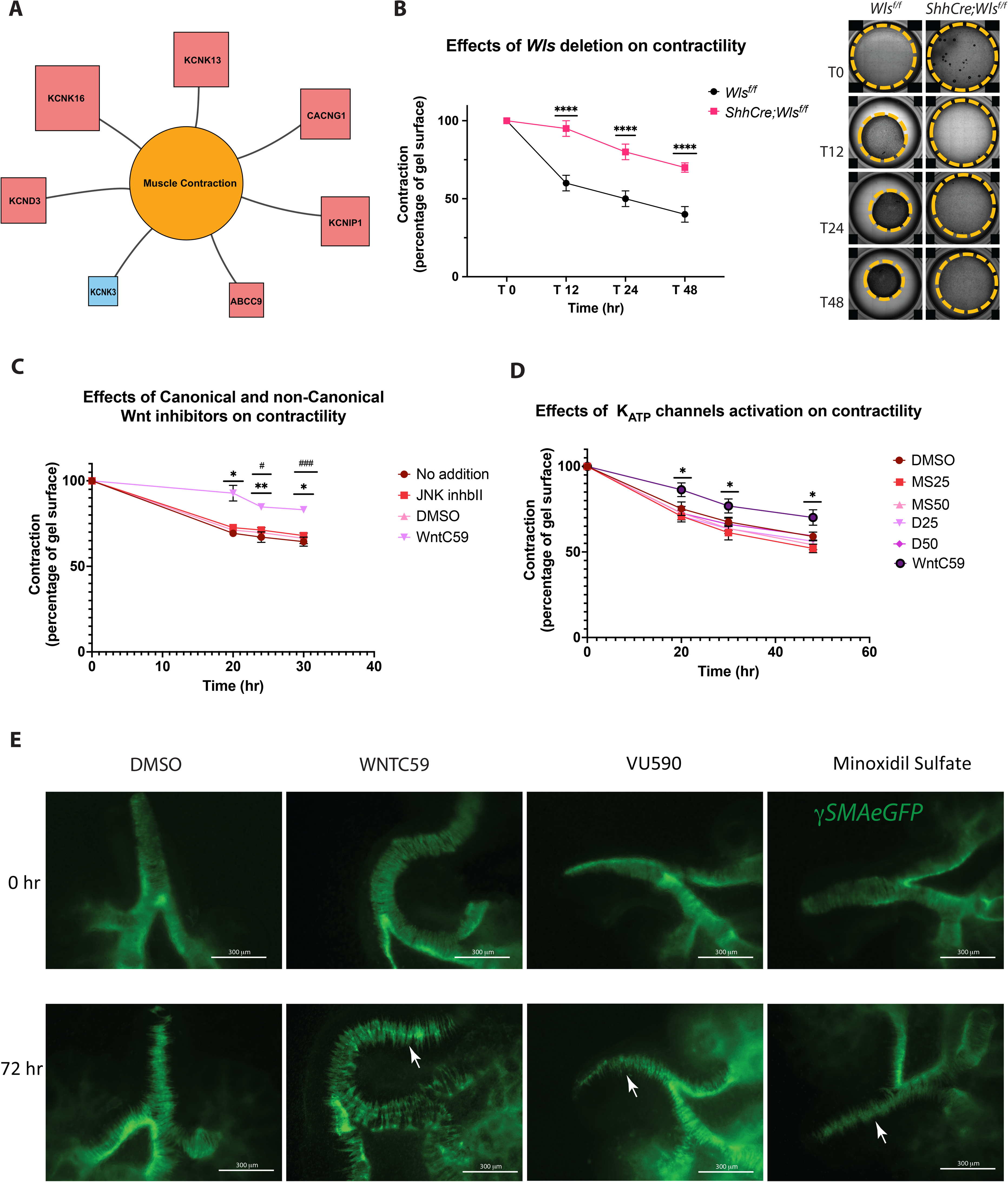
Abnormal ion channel activity and deficient Wnt signaling impair the organization and contractility of the developing trachealis muscle ex vivo. A) Wls-regulated ion channels influence muscle contractility. Diagram depicting functional relationships of differentially regulated ion channels on muscle contractility was generated using RNA seq data from E13.5 *ShhCre;Wls^f/f^*tracheas. Red boxes indicate genes which are upregulated, and blue boxes indicate those which are downregulated. The size of the boxes represents fold change expression after epithelial deletion of *Wls*. B) Deletion of *Wls* impairs contractility of tracheal mesenchymal cells in free floating collagen assay. Bright field images depict the extent of the contraction in *Wls^f/f^*primary cells determined by the reduction in the size of the surface of the collagen gel. *ShhCre;Wls^f/f^* cells did not contract and the gel surface remained almost unchanged throughout the experiment. Graph denotes contractility of primary tracheal mesenchymal cells determined by measurement of collagen gel surface T-test ****p<0.0005 N=7. C) Inhibition of Wnt signaling by pharmacological treatment with WntC59 impaired contractility of cells. No significant effect on contractility was observed when Wnt/β-catenin independent signaling was pharmacologically inhibited with JNK inhibitor II. Two-way ANOVA, *p< 0.05 WntC59 vs No addition, JNKinhII, DMSO, **p<0.01vs WntC59 vs No Addition, DMSO, #p<0.05 vs JNK inhII, ###p<0.001 WntC59 vs JNK inhII N=3 D) Activation of Kcnj8 channel with Minoxidil or Diazoxide, did not affect the contractility of primary tracheal mesenchymal cells, while WntC59 impaired contraction. Two-way ANOVA *p<0.05 WntC59 vs DMSO, MS25 (Minoxidil Sulfate 25uM), MS50 (Minoxidil Sulfate 50uM), D25 (Diazoxide 25uM), D50 (Diazoxide 50uM) N=4. E) Trachea lung explants isolated at E12.5 were cultured in the air-liquid-interface (ALI) over 72 hours. Tissue treated with WntC59 or VU590, displayed disorganized trachealis smooth muscle cells lacking cell-cell adhesion (arrows) and disrupted cell orientation. Minoxidil sulfate treatment did not alter smooth muscle organization while compared to tissue treated with DMSO. Representative images are shown.

### Mesenchymal condensation is altered after inhibition or activation of inwardly rectifying ion channels

Using a mouse model wherein Sox9+ cells (chondroblasts) are genetically labeled with GFP we performed ALI cultures of tracheal-lung explants isolated at E11.5 for 72 hrs to observe the process of mesenchymal condensations that occurs *in vivo* around E12.5 and is completed by E14.5. The formation of mesenchymal condensations (arrows) necessary for cartilage formation was severely impaired after pharmacological inhibition of Wnt signaling with WntC59 or potassium ion channel activity with VU590 (Fig5A). On the other hand, activation of the potassium ion channel Kcnj8 with minoxidil sulfate delayed and impaired the mesenchymal condensation but did not prevent the formation of cartilage (Fig5B). Further, using the micromasses system, we tested whether activity of the KCNJ ion channels (KCNJ13 and KCNJ8) affects the formation of the extracellular matrix necessary for cartilage formation. In general, when micromasses were seeded and treated with VU590, minoxidil sulfate or diazoxide in conjunction with BMP4, formation of cartilaginous extracellular matrix (ECM) was severely impaired. In contrast, treatment with BMP4 only resulted in deposition of cartilaginous ECM as determined by alcian blue staining (Fig5C). Next, we tested if activity of the ion channels was required in cells before the process of the mesenchymal condensation. Mesenchymal cells were pretreated in monolayers with VU590, or minoxidil sulfate, seeded in micromasses, and treated with BMP4. While cells pretreated with minoxidil sulfate condensed and secreted ECM to some extent, cells pretreated with VU590 did not produce ECM. Thus, KCNJ13 may play a critical role before and during chondroblast condensation, while enhanced KCNJ8 activity appears to affect later stages of chondrogenesis once cells are already condensed. Taken together inwardly rectifying ion channels play a role in mesenchymal condensations and cartilage formation.

**Figure5:**
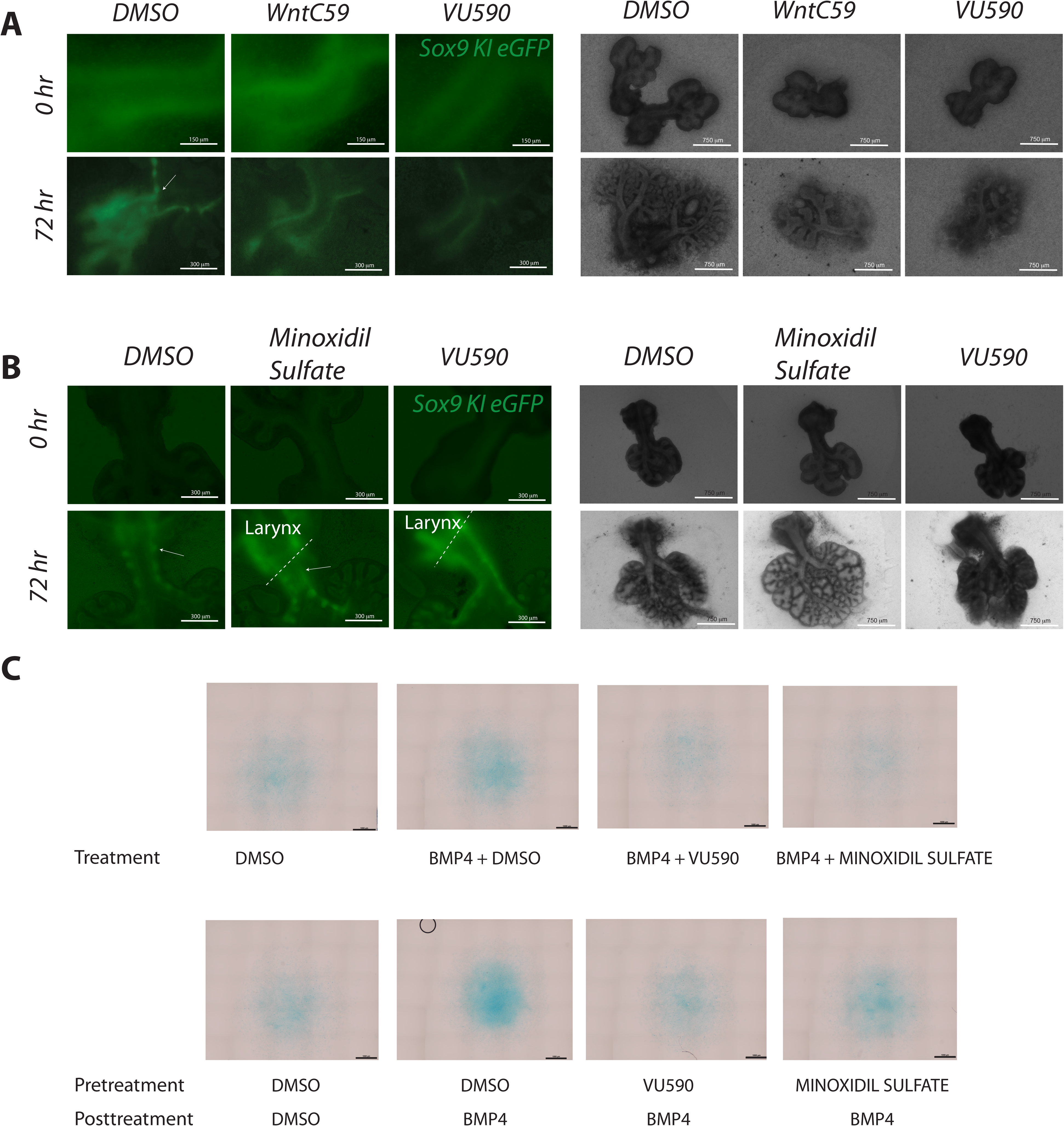
Mesenchymal condensation is altered by inhibition or activation of inwardly rectifying ion channels. A) Trachea-lung explants isolated from E12.5 *Sox9KIeGFP* embryos were cultured in ALI for 72 hours. Mesenchymal condensations required for cartilage formation were present in the DMSO tissue after 72 hr incubation (arrow). Inhibition of Wnt signaling with WntC59 or inhibition of potassium ion channel with VU590 blocked mesenchymal condensations. Corresponding low magnification bright field images of the cultures are shown to the right of fluorescent images. B) Activation of K_ATP_ ion channels delayed mesenchymal condensations without substantially impacting the growth of the trachea and lung branching as determined by bright field images. C) Primary tracheal mesenchymal cells were seeded at high concentration in micromasses. Treatment with rBMP4 produced extracellular matrix as determined by the Alcian blue staining (compare to DMSO). Combined treatment with rBMP4 and VU590 or rBMP4 and Minoxidil sulfate severely impaired the secretion of extracellular matrix as determined by the lack of Alcian blue staining. Pretreatment of primary tracheal mesenchymal cells with VU590 before seeding in micromasses prevented secretion of cartilaginous extracellular matrix as determined by the lack of Alcian blue staining in micromasses. Pretreatment with Minoxidil sulfate, an activator of K_ATP_ channels, did not significantly affect production of extracellular matrix in micromasses.

## DISCUSSION

In the present study, we focused on mechanisms by which epithelial-induced Wnt signaling influences trachealis muscle organization and patterning. In developing tracheal mesenchyme, Wnt/β-catenin signaling downstream of Wls is essential for the patterning and assembly of smooth muscle cells. Further, Wnt/β-catenin regulates the expression of genes encoding potassium ion channels, including the inwardly rectifying ion channels *Kcnj13* and *Kcnj8* and the voltage-gated channel *Kcnd3*. Remarkably, the role of β-catenin is in repressing the expression of ion channels, thus modulating the activity of the channels necessary for muscle cell assembly and cartilaginous mesenchymal condensations during tracheal development (Fig. 6).

**Figure 6:**
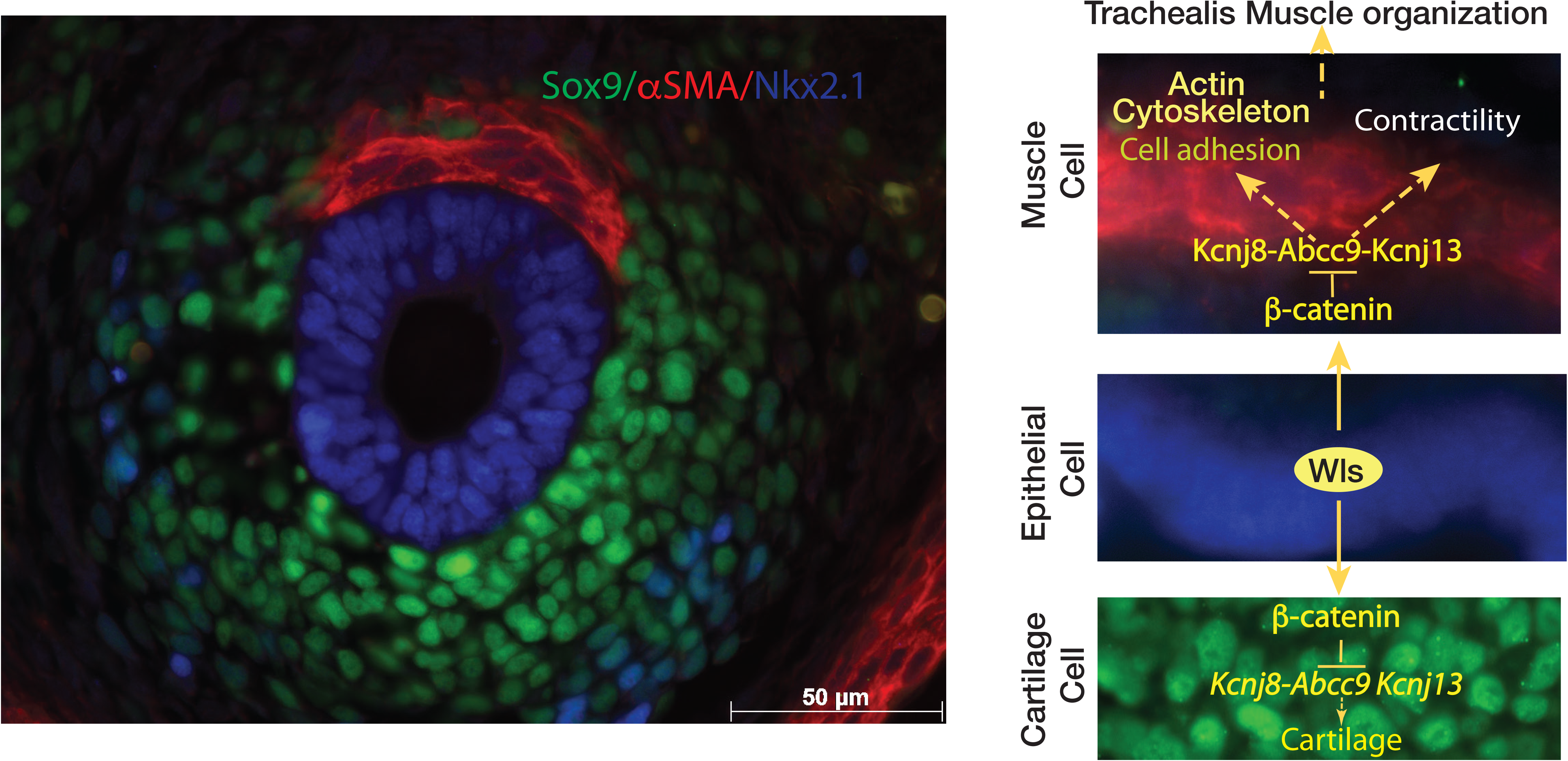
Model. Transverse section of a E13.5 mouse trachea depicting trachealis smooth muscle (αSMA) and ventrolateral cartilage (Sox9) is shown. Epithelial Wls-induced Wnt signaling, modulates expression of ion channels including *Kcnj8-Abcc9* and *Kcnj13*. We propose that ion channels facilitate the rearrangement of the smooth muscle cell fibers, smooth muscle cell contractility and adhesion, and influences cartilage formation.

### Unique role for Wnt/β-catenin and non-canonical Wnt signaling in trachealis smooth muscle patterning and organization

In developing trachea, Wnt signaling differentially regulates expression of genes promoting the organization of the actin cytoskeleton as determined by unbiased analysis of the gene expression in *Wls* deficient trachea. These data are supported by the morphological changes observed in the trachealis muscle of the *ShhCre;Wls^f/f^*embryos, wherein anomalous muscle cell shape was detected (Fig.1).

Wnt/β-catenin independent signaling has been shown to play a critical role in cell polarization and behavior during development and disease (Arabzadeh et al. 2016; Cheng et al. 2008; De Calisto et al. 2005; Gong, Mo, and Fraser 2004). Wnt/β-catenin independent signaling is also critical for the proper assembly and polarization of trachealis smooth muscle cells taking place between E12.5 and E14.5, a period when cartilaginous mesenchymal condensations are occurring (Hines et al. 2013). Deletion of *Wnt5a* and *Ror2*, targets of epithelial Wls-induced-Wnt signaling (Sinner et al. 2019), impair the polarization and contractility of the trachealis muscle affecting the tracheal tube elongation (Oishi et al. 2003; Kishimoto et al. 2018). Although smooth muscle cell organization and assembly were affected after deletion of *Wnt5a*, dorsal-ventral patterning remained unaltered as the smooth muscle cells remained dorsally positioned (Fig1,3). Thus, noncanonical Wnt signaling mediated by Wnt5a is essential for polarization of the trachealis muscle but dispensable for dorsal-ventral patterning mediated by epithelial Wnt ligands.

We also observed that at E13.5 Wnt4, a ligand also associated with β-catenin independent Wnt signaling(Louis et al. 2008; Tanigawa et al. 2011), was detected in the dorsal aspect of the epithelium and to lesser degree in the mesenchyme of the developing trachea. Previous studies have demonstrated a role of Wnt4 in pulmonary growth and tracheal cartilage formation by modulation of Sox9 expression(Caprioli et al. 2015); however, it is unknown whether Wnt4 may also impact the organization of the trachealis muscle. Further studies are required to determine its role in smooth muscle differentiation.

Our studies determine that mesenchymal Wnt/β-catenin activity in developing trachea is required for patterning and organization of trachealis muscle. Wnt/β-catenin and its direct targets are reduced in the tracheal mesenchyme of *Wls* deficient trachea (Bottasso-Arias et al. 2022; Gerhardt et al. 2018; Snowball et al. 2015). After mesenchymal deletion of *β-catenin*, cells giving rise to cartilage are drastically reduced while smooth muscle is increased, recapitulating the phenotype seen after epithelial deletion of *Wls* (Fig2). These data are supported by other studies, confirming the role of β-catenin in tracheal cell specification (Kishimoto et al. 2020; Hou et al. 2019). Remarkably we also found that besides the ectopic localization of the trachealis smooth muscle cells, myocytes were poorly organized. The lack of organization of the muscle cells suggests a role for β-catenin in cell polarization, likely by induction of genes promoting cytoskeletal organization.

While deletion of Wls has a marked effect in tracheal mesenchymal cell differentiation and cytoskeletal organization, we did not observe significant morphological changes in *Wls* deficient epithelial cells (SFig1). Moreover, at the stages analyzed epithelial cells possess a clear respiratory identity as determined by *Nkx2.1* expression (Fig1). However, studies performed at later developmental stages demonstrated that epithelial deletion of *Wls* causes a significant reduction in the number of basal progenitor cells of the tracheal epithelium (Hou et al. 2019).

### Wnt/β-catenin as a modulator of ion channel expression

Unbiased analysis of gene expression determined that Wnt signaling influences expression of ion channels in developing trachea. Among the ion channels differentially regulated we identified ion channels controlling potassium balance (Fig.3 A, B). Potassium channels regulate the membrane potential of smooth muscle cells, which in turn regulates the cytoplasmic free Ca^2+^ concentration. Calcium levels are critical for cell signaling activity as well as cell-cell interactions (S.A. Kim et al. 2011). Potassium channels also affect the actin cytoskeleton organization of developing trachealis smooth muscle (Yin et al. 2018). Therefore, changes observed in genes associated with the actin cytoskeleton and ion channels are influenced by Wnt signaling during a critical stage of tracheal development, when mesenchymal cells differentiate and organize to give rise to muscle and cartilage (Hines et al. 2013; Kishimoto et al. 2018).

One set of differentially regulated genes includes potassium-voltage gated channel encoding genes. KCNC2 (Kv3.2) (Haas et al. 1993) is expressed in cells destined to give rise to tracheal cartilage. *KCNC2* mutations are associated with developmental delay, ataxia and abnormal metabolism linked to type II diabetes mellitus (Hwang et al. 2016). A variant for *KCNC2* was identified by GWAS analysis among smokers, suggesting a role for KCNC2 in pulmonary function (Hilger et al. 2015). Further, *KCNC2* is among the signature genes detected in lymphangioleiomyomatosis (LAM) core cells of the diseased lung (Guo et al. 2020). At present, KCNC2’s role in tracheal cartilage development remains to be determined. *Kcnd3* is expressed in tracheal smooth muscle cells and its expression was increased after deletion of *Wls* or β-catenin (Fig2,3). Mutations in *KCND3* have been associated with Brugada Syndrome, Spinocerebellar Ataxia and Atrial fibrillation (Hsiao et al. 2021; Olesen et al. 2013). A recent study on consensus gene co-expression network analysis identified *KCND3* as a novel gene associated with the severity of idiopathic pulmonary fibrosis (Ghandikota et al. 2022). It is unclear whether *KCND3* plays a role in tracheal development; however, transcriptomic analysis of fetal tracheoesophageal human tissue detected overexpression of *KCND3* in tracheoesophageal fistula tissues (Brosens et al. 2020).

Another set of differentially regulated ion channels includes the inwardly rectifying ion channel KCNJ8 and its regulatory subunit ABCC9. *Kcnj8* and *Abcc9* are expressed in the lung and may play a role in wound healing by promoting migration via Erk1/2 activation (Buchanan et al. 2013). Based on our studies *Kcnj8, and Abcc9* RNA are ectopically expressed throughout the trachea after deletion of *Wls* (Fig.2C). Further, *Kcnj8* and *Abcc9* are upregulated after mesenchymal deletion of *β-catenin*, but not after mesenchymal deletion of *Wnt5a* (Fig.3C).

### Channelopathies and structural anomalies of the upper airways

A growing body of research indicates that channelopathies are underlying causes of disease for pathologies occurring in tissues and systems beyond the muscle and nervous system (J.B. Kim 2014). In the respiratory tract, the main pathology observed associated with abnormal ion channel function or expression is cystic fibrosis. The pathology is the result of mutated *CFTR*, encoding a chlorine channel located in the apical membrane of the epithelium. Patients with cystic fibrosis are vulnerable to severe and chronic pulmonary infections and inflammation, which lead to irreversible airway damage and respiratory failure in most cases. While some prevalence of tracheomalacia has been described in infants diagnosed with cystic fibrosis, it is unclear if the abnormal function of the channel is responsible for the structural defect on the central airway or it is secondary to infections causing damage to the airways(Fischer et al. 2014). However, in mouse models, deletion of *Cftr* results in abnormal patterning of the trachea and reduction of the luminal surface (Bonvin et al. 2008; Meyerholz et al. 2010). Other genes encoding ion channels associated with tracheal defects in mouse include *Kcnj13, Tmem16a and Cav3.2* (Yin et al. 2018; Rock, Futtner, and Harfe 2008; Almeida et al. 2007). In our studies, we identified *Cftr, Kcnj13, and Tmem16a* as genes differentially regulated by Wnt/β-catenin signaling in a model of tracheomalacia, supporting a role for the genes in central airway patterning.

In humans, mutations in *KCNJ8* and *ABCC9* underlie the pathology of Cantu syndrome (Cooper et al. 2014). The pathology is associated with an array of defects, and the presentation is variable. Activating mutations in *KCNJ8* are associated with respiratory distress and malacias of the upper airways and bronchi (Grange et al. 2019). Kcnj8 was differentially regulated and based on our genetic analysis, Kcnj8 is repressed by Wnt/β-catenin signaling during tracheal development (Fig3). Further, pharmacological treatment targeting Kcnj8 affected chondrogenesis in ex vivo system (Fig5). While a direct relationship between *KCNJ8* and tracheal cartilage has not been established, we identified rare heterozygous variants for *ABCC9*, the associated gating subunit of *KCNJ8*, and other potassium ion channels in exome sequencing data obtained from CTR trios (Sinner et al. 2019). However, further examination and testing are required to determine whether the variants could have an impact in cartilage development.

### Final Remarks

A better understanding of the mechanisms mediating the formation of large airways can provide valuable information on the molecular mechanisms of pathologies such as TBM and CTR and, thus, contribute to improve diagnosis and treatments. The present work highlights how the Wnt signaling pathway influences ion channel expression that is necessary for proper tracheal formation. Given their impact on tracheal development, ion channels are good candidates for therapeutic drugs currently available that may be repurposed to treat TBM or CTR.

## Supporting information

Control video still image

Mutant video still image

Mutant trachea video

Control trachea video

## Acknowledgements

We appreciate the insightful comments on the manuscript by Dr. Whitsett and Dr. Swarr. We are grateful to Chuck Crimmel for assistance with graphic design. This work was partially supported by National Institutes of Health NHLBI R01 144774 to DS. MSK was supported by the National Institutes of Health (1R01 HL134801 and 1R01 HL157176).

