## Supplementary figures and images for "Wnt signaling regulates ion channel expression to promote smooth muscle and cartilage formation in developing mouse trachea"

### Control video still image

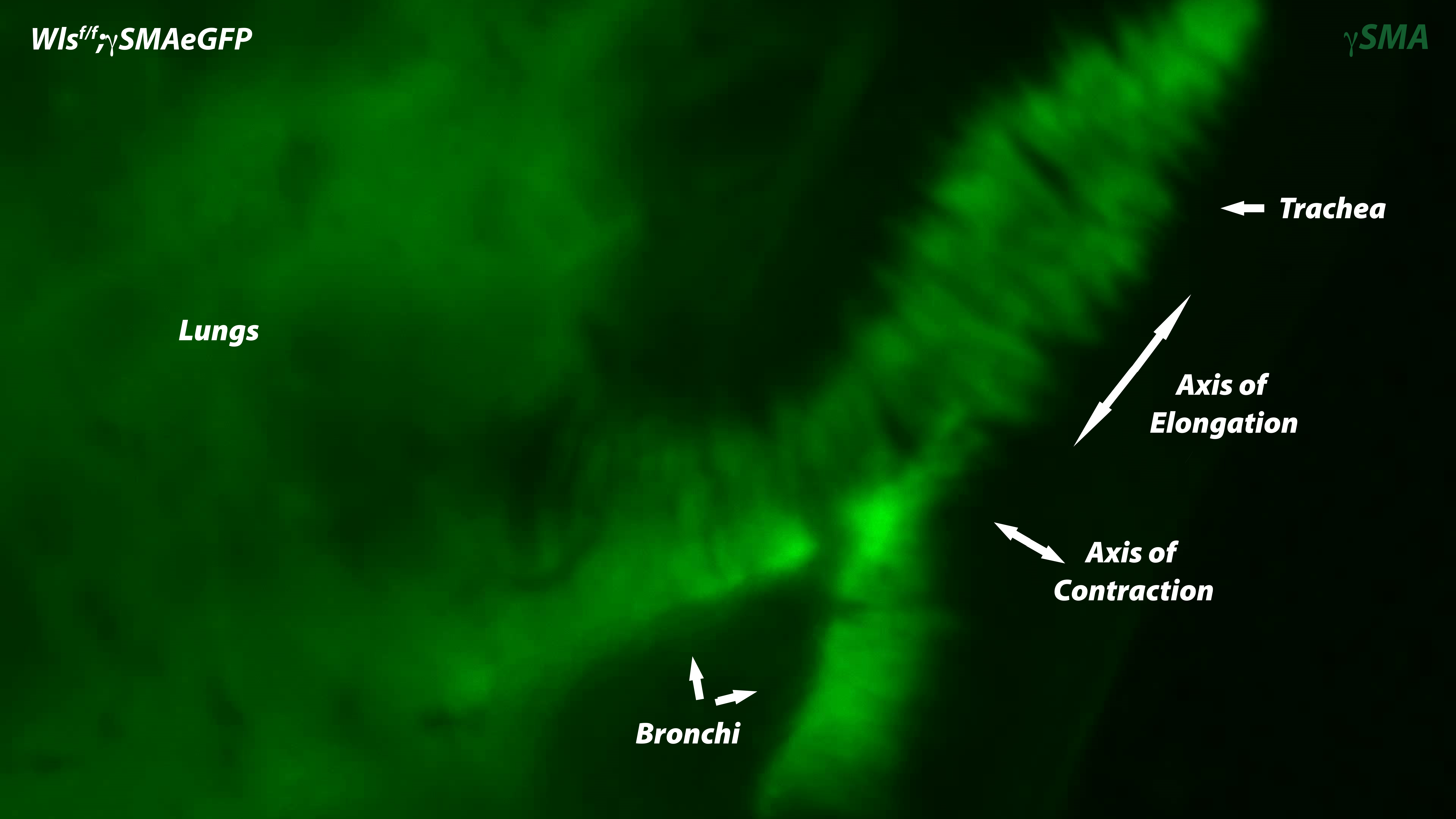

### Mutant video still image

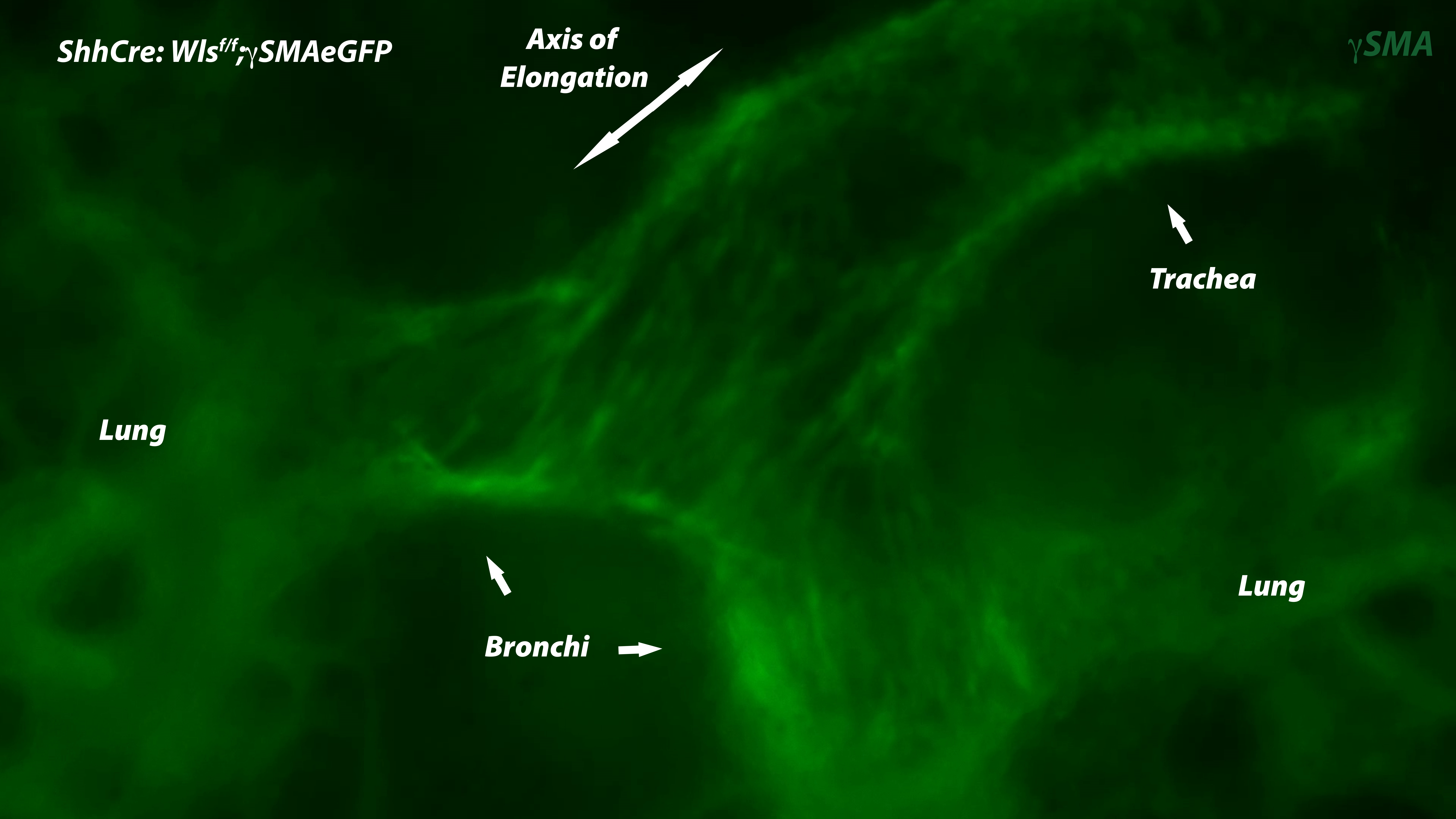

### Supplementary figure 1

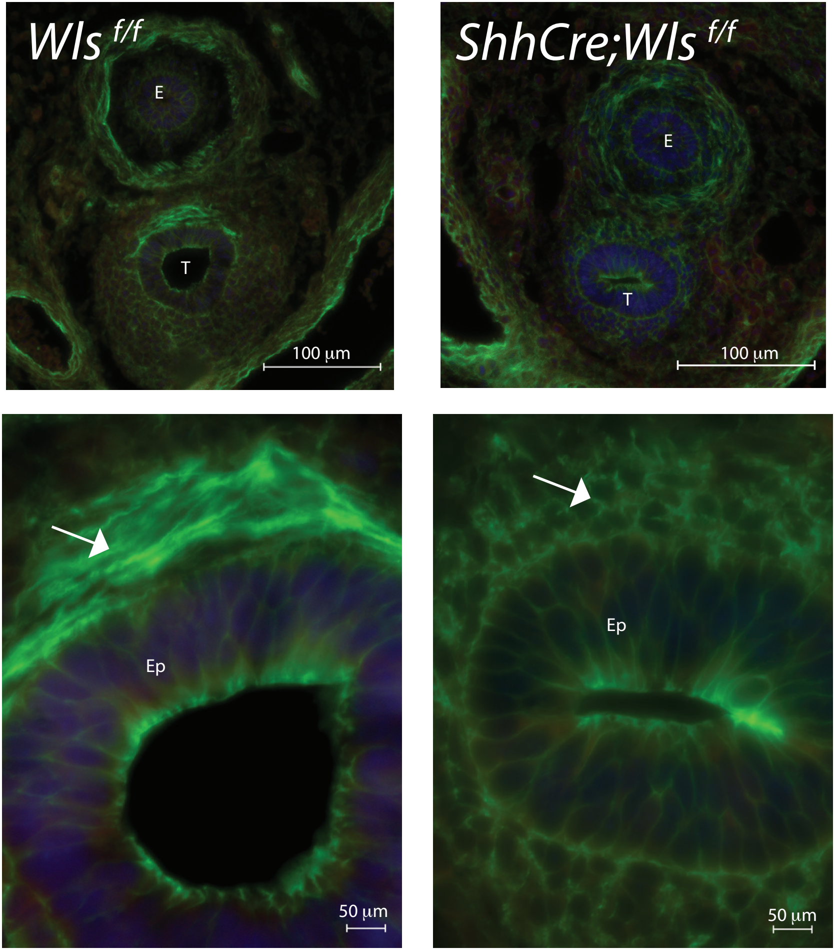

### Supplementary Figure 2

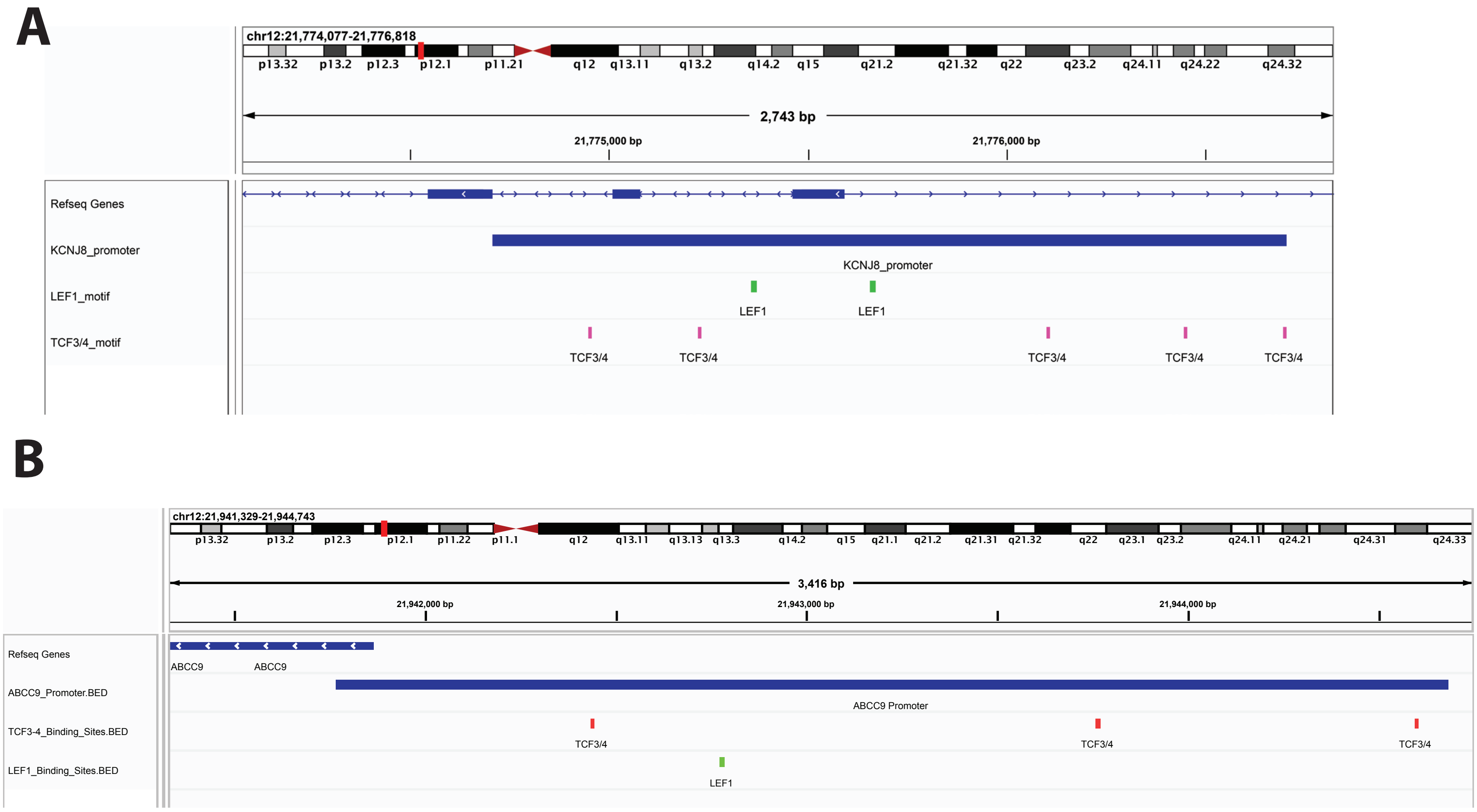

### Supplementary Figure 3

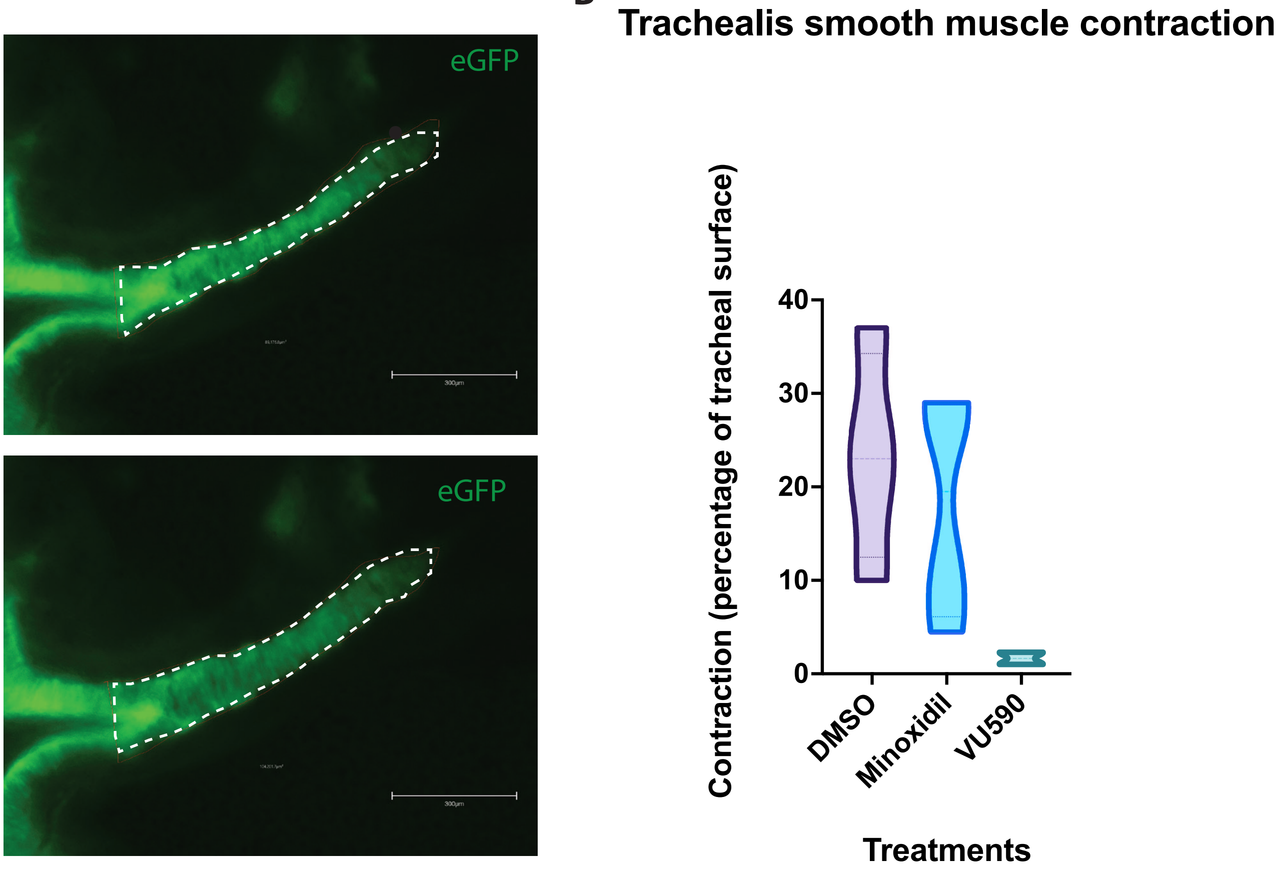
