## Supplement information for "Wnt signaling regulates ion channel expression to promote smooth muscle and cartilage formation in developing mouse trachea"

**SUPPLEMENTARY MATERIAL**

**Table1:**

Primers utilized for genotyping.

**Table 2:**

Antibodies used for the studies.

**Figures**

**Fig S1:**

Phalloidin staining on transverse sections demonstrates the organization of actin filaments in E13.5 control and *ShhCre; Wls^f/f^ tracheal* and esophageal tissue. Note the circular shape in smooth muscle cells of the mutant trachea (arrows in high magnification). No differences in cell shape were detected between the control and mutant epithelial cells. E= esophago, T=trachea, ep: epithelium.

**Fig S2:**

Lef1 and TCF3/4 binding sites are present in *KCNJ8* and *ABCC9* promoters.

In silico analysis of human *KCNJ8* and *ABCC9* promoters, identified putative binding sites for LEF1 and TCF3/4 in a 2KB region proximal to the Transcriptional Starting Site.

**Fig S3:**

Effect of pharmacological activation or inhibition of ion channels on trachealis smooth muscle cell contraction *ex vivo*. *γSMAeGFP* trachea-lung tissue were cultured in ALI and treated with DMSO, Minoxidil sulfate or VU590. After 48 hours, thirty-second videos were recorded. Surface of the trachealis muscle was measured at the peak of contraction and when the cells were relaxed determining the amplitude of the contraction. A) Examples of contracted (top immunofluorescence image) or relaxed (bottom image) trachealis muscle is shown. B) Contraction was not affected by treatment with Minoxidil while VU590 treatment prevented contraction in *ex vivo* assay.

**Video:**

Trachealis muscle contraction of *Wls^f/f^;γSMAeGFP* (video1) and *ShhCre; Wls^f/f^;γSMAeGFP* (video2) were captured during thirty-second videos. Note the orientation of the contraction is transversal to the elongation axis of the trachea in control *Wls^f/f^;γSMAeGFP,* which is not observed in the mutant trachea *ShhCre; Wls^f/f^;γSMAeGFP*. However, contraction was observed in the bronchi of the *ShhCre; Wls^f/f^;γSMAeGFP*.
