## Supplement Table 1 for "Wnt signaling regulates ion channel expression to promote smooth muscle and cartilage formation in developing mouse trachea"

PRIMERS FOR GENOTYPING:

Wls F- 5’ AGG CTT CGA ACG TAA CTG ACC 3’
Wls R- 5’ CTC AGA ACT CCC TTC TTG AAG C 3’

Shhcre 1- 5’ GGG ACA GCT CAC AAG TCC TC 3’
Shhcre 2- 5’ CTC GGC TAC GTT GGG AAT AA 3’
Shhcre 3- 5’ GGT GCG CTC CTG GAC GTA 3’

Dermo1Cre/Foxg1cre F- 5’ TGC CAC GAC CAA GTG ACA GCA ATG 3’
Dermo1Cre/Foxg1cre R- 5’ AGA GAC GGA AAT CCA TCG CTC G 3’

Wnt5a F- 5’ GGT GAG GGA CTG GAA GTT GC 3’
Wnt5a R- 5’ GGA GCA GAT GTT TAT TGC CTT C 3’

β-Catenin Floxed F- 5’AAG GTA GAG TGA TGA AAG TTG TTG TT 3’
β-catenin Floxed R- 5’ CAC CAT GTC CTC TGT CTA TTC 3’

Sox9 KI 1- 5’ GAG GGG CTT GTC TCC TTC AG 3’
Sox9 KI 2- 5’ ACA CCG GCC TTA TTC CAA G 3’
Sox9 KI 3- 5’ GGC AGC TAC TCT TGA AAT CCA 3’

Ror2 F- 5’ TGC AGG TTT TGA GCC CTA AC 3’
Ror2 R- 5’ CGA GAA TGA CTT CCC TGT CC 3’

γSMAeGFP F- 5’ CCT ACG GCG TGC AGT GCT TCA GC 3’
γSMAeGFP R- 5’ CGC CGA GCT GCA CGC TGC GTC CTC 3’
