## Supplement table 2 for "Wnt signaling regulates ion channel expression to promote smooth muscle and cartilage formation in developing mouse trachea"

| PRIMARY ANTIBODY | REFERENCES | SPECIES | DILUTION | COMPANY | CATALOG # |
|---|---|---|---|---|---|
| Anti Actin, Alpha Smooth Muscle-Cy3, monoclonal | [1] | Mouse | 1:200 | Sigma Aldrich | C6198 |
| Anti Actin, Alpha Smooth Muscle, monoclonal | [2] | Mouse | 1:200 | Sigma Aldrich | A5228 |
| Anti-Sox9, polyclonal | [2] | Rabbit | 1:200 | Millipore | AB5535 |
| Human Sox9, polyclonal | [3] | Goat | 1:50 | R&D | AF3075 |
| Nkx2.1 | [4] | Rabbit | 1:200 | Seven Hills Bioreagents | R1231 |
| Nkx2.1 | [5] | Guinea Pig | 1:200 | Seven Hills Bioreagents | GP237 |
| **SECONDARY ANTIBODY** | | | | | |
| Donkey anti Mouse IgG (H+L), 350 | | | 1:200 | Invitrogen | A10035 |
| Donkey anti Rabbit IgG (H+L), 350 | | | 1:200 | Invitrogen | A10039 |
| Donkey anti Goat IgG (H+L), 488 | | | 1:200 | Jackson Immuno | 705-546-147 |
| Donkey anti Guinea Pig (H+L), 488 | | | 1:200 | Jackson Immuno | 706-545-148 |
| Donkey anti Mouse IgG (H+L), 488 | | | 1:200 | Invitrogen | A21202 |
| Donkey anti Rabbit IgG (H+L), 488 | | | 1:200 | Invitrogen | A21206 |
| Donkey anti Mouse IgG (H+L), 594 | | | 1:200 | Invitrogen | A21203 |
| Donkey anti Rabbit IgG (H+L), 594 | | | 1:200 | Invitrogen | A21207 |
